# A simplified High-Pressure Freezing Workflow in Autogrids

**DOI:** 10.64898/2026.04.18.719352

**Authors:** Miriam S. Weber, Till Epprecht, Jingwei Xu, Gregor L. Weiss

**Affiliations:** Institute of Medical Microbiology, University of Zürich, 8006 Zürich, Switzerland; Westlake Laboratory of Life Sciences and Biomedicine, Hangzhou, 310024, Zhejiang, China; School of Life Sciences, Westlake University, Hangzhou, Zhejiang 310024, China

**Keywords:** High-Pressure Freezing, plasma-focused ion beam milling, Waffle method, on-grid lamella preparation, cryo-lift out, SOLIST, cryo-electron tomography, subtomogram averaging

## Abstract

Cryo-electron tomography (cryo-ET) is a powerful method for studying biological structures in near-native states, yet vitrification of thick and complex specimens remains a major limitation. High-pressure freezing (HPF) allows vitrification of up to ∼200 µm thick samples, but its seamless integration into cryo-electron microscopy (cryo-EM) workflows is hindered by planchettes that are not optimized for EM grids, resulting in extensive handling, reliance on user experience, and frequent grid damage. Here, we present a streamlined HPF workflow based on newly designed planchettes that directly accommodate clipped EM Autogrids. Combined with interchangeable lids for controlling sample thickness, this system enables robust vitrification of diverse specimens, including single-celled eukaryotes and tissue biopsies, while minimizing post-freezing handling. Integration with plasma-focused ion beam milling supports both on-grid lamella preparation and lift-out approaches. We demonstrate reliable preservation of cellular ultrastructure and macromolecular integrity by cryo-ET and subtomogram averaging. This resource provides a practical and scalable solution for integrating HPF into modern cryo-ET pipelines, expanding access to structurally complex biological systems and supporting future clinical applications.

## Introduction

Cryo-electron tomography (cryo-ET) provides near-native structural insights by imaging proteins and organelles directly in their cellular environment, without the need for protein purification, chemical fixation or heavy metal staining. This approach enables visualization of cellular processes and interactions at sub-nanometer resolution and in a fully hydrated state ^1^. For thin biological specimens of up to ∼10 µm in thickness, such as bacteria and individual eukaryotic cells, plunge freezing is the method of choice for cryo-fixation and reliably preserves native ultrastructure ^2^. While revolutionary discoveries have been made over the past decades, the field is increasingly shifting toward studying more complex biological structures and organisms ^3–10^, enabled by constant developments in instrumentation and software ^11–16^. Many of the most promising frontiers in today’s cellular structural biology – including the direct analysis of environmental samples, bacterial biofilms, multicellular organisms, 3D organoids, and tissue or patient biopsies – exceed the thickness limit of plunge freezing and remain challenging to study ^17–19^. Addressing these samples requires alternative vitrification methods combined with advanced sample-thinning strategies.

High-pressure freezing (HPF), developed more than 50 years ago, is currently the only technique capable of vitrifying biological material up to ∼200 µm in thickness ^20–25^ and is routinely used in combination with freeze-substitution, resin embedding and conventional electron microscopy workflows such as focused ion beam (FIB)-scanning electron microscopy (SEM). For HPF, samples are placed between metal carriers, so-called planchettes, and rapidly vitrified within milliseconds by the application of pressurized liquid nitrogen. The advent of advanced sample thinning techniques has renewed interest in HPF as an entry point for structural studies of complex multicellular systems using cryo-ET ^4,8,10,26,27^. Approaches such as the Waffle method for freezing directly on EM grids ^28^, or downstream processing strategies like cryo-EM of vitreous sections (CEMOVIS) ^29,30^ as well as on-grid lamella preparation ^7,28,31,32^ and lift-out approaches ^3,10,26,27,33,34^ using cryo-FIB instruments have extended its applicability. Despite its potential though, HPF itself has seen little technical innovation and remains difficult to integrate into routine cryo-EM workflows.

A key bottleneck for HPF-based cryo-EM analysis is sample handling: specimens must either be recovered from the planchette post-freezing by complex lift-out strategies or be frozen directly on fragile, unclipped EM grids for on-grid lamella preparation. Conventional HPF planchettes are typically available in 3 mm or 6 mm diameters and have either flat surfaces or spacer cavities of variable depth to accommodate different sample types. However, neither format is well suited for freezing samples directly on EM grids. The use of 3 mm planchettes requires adaptations of HPF equipment, since they do not tolerate the slightly larger EM grids (diameter of 3.05 mm). In contrast, 6 mm planchettes provide excess space, require larger sample volumes, and do not constrain specimen movement. Importantly, conventional planchettes lack a dedicated grid-holding geometry, allowing grids to shift during preparation and freezing, which is particularly problematic for tissue samples. After freezing, grids frequently adhere to the metal surface of the planchette and are prone to bending and damage during retrieval. In contrast, cryo-FIB workflows have transitioned to using EM grids that were clipped into rigid metal support rings called Autogrids ^12,35^, which improve mechanical stability, reduce handling damage, and are compatible with modern autoloader systems in plasma-FIB (pFIB) instruments and transmission electron microscopes (TEM). Direct freezing in Autogrids has recently been introduced in jet-freezing devices ^36^ and even for plunge freezing approaches ^37^. However, traditional HPF planchettes are incompatible with clipped EM grids, resulting in a largely manual workflow that requires substantial user expertise, and thereby limit broader adoption of HPF.

To overcome these limitations and to facilitate broader use of HPF in cryo-EM workflows, we developed and benchmarked a new generation of planchettes specifically designed to accommodate clipped EM Autogrids. This design enables stable grid positioning during freezing, prevents deformation, and minimizes post-freezing handling. We demonstrate the robustness of this system across multiple sample types and establish improved cryo-FIB milling approaches to validate structural preservation by analyzing cellular architecture by cryo-ET and subtomogram averaging.

## Results

### Designing Autogrid Planchettes for streamlined High-Pressure Freezing

To enable direct HPF on clipped Autogrids, we developed a new Autogrid planchette system comprising two components: a bottom planchette that holds the Autogrid, and interchangeable lids tailored to different sample types and thicknesses.

The Autogrid planchette features a dedicated grid-holding geometry that positions the commercially available Autogrids precisely during freezing **(Figure 1A, B)**. A circular cutout secures the Autogrid, while a raised central platform supports the grid mesh and prevents deformation during freezing. Additional side cutouts enable retrieval of the frozen grid with tweezers, minimizing mechanical damage during extraction **(Figure 1A, B)**.

**Figure 1.**
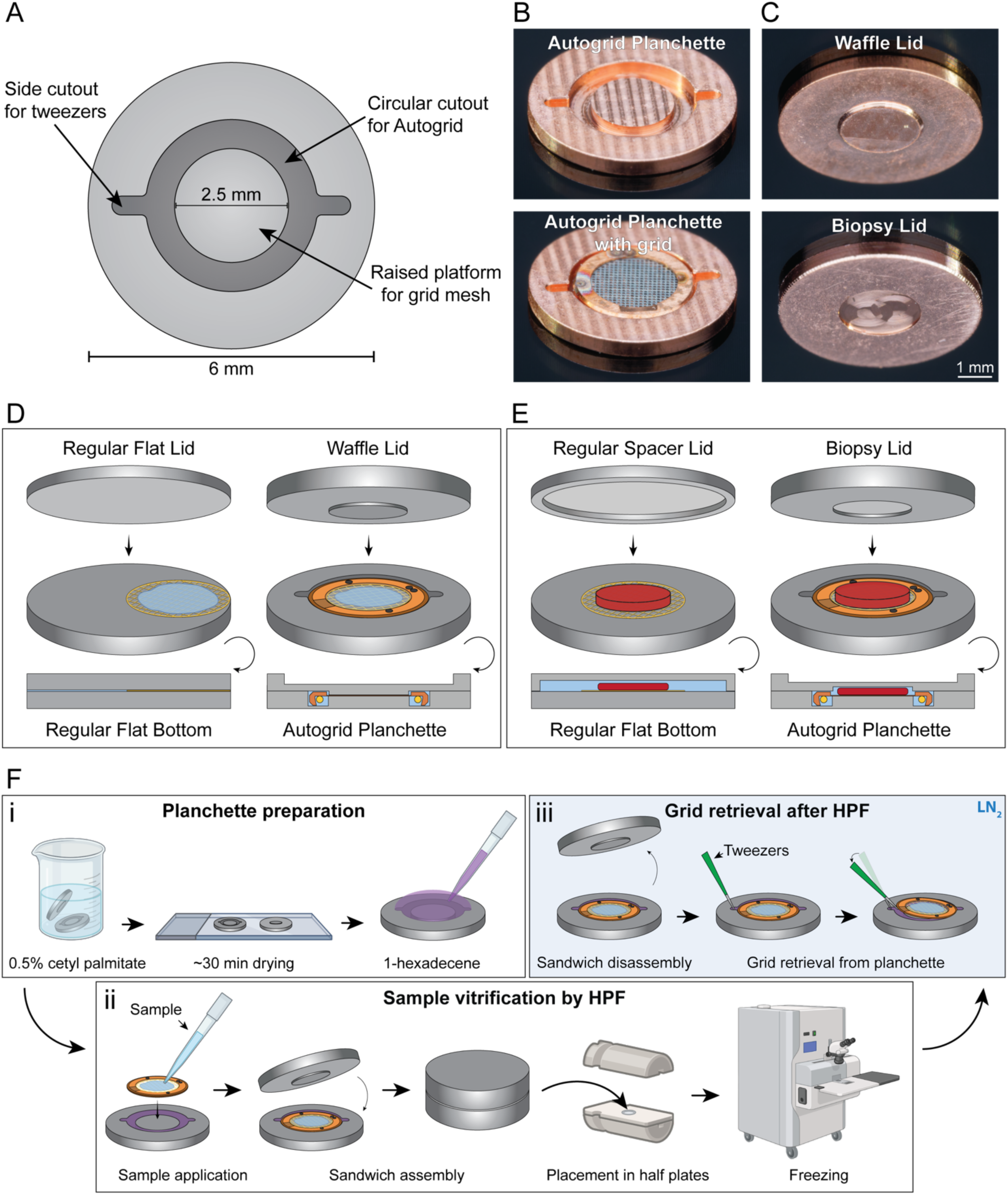
Designing Autogrid planchettes for streamlined high-pressure freezing. **(A)** Schematic of the Autogrid planchette designed to accommodate EM grids clipped into Autogrids. **(B)** Representative photographs of a copper Autogrid planchette with and without clipped EM-Autogrid. **(C)** Representative photographs of copper Waffle and Biopsy lid planchettes. **(D)** Schematic comparison of conventional HPF planchettes and the Autogrid planchette system for Waffle-style freezing. In conventional workflows, flat planchettes are used on either side of the unclipped grid. In contrast, the Autogrid planchette system uses a bottom planchette with a dedicated grid holding geometry to position the clipped grid, combined with a lid that seals on top of the grid mesh to confine the sample between grid bars. Cut-open side views illustrate the positioning of the Autogrid within the planchette and the tight sealing of the lid. **(E)** Schematic comparison of conventional HPF planchettes and the Autogrid planchette system for freezing tissue biopsies. Conventional carriers require filling a large volume surrounding the grid with buffer, whereas the Biopsy lid restricts the sample and buffer to the grid mesh area, excluding the Autogrid ring and surrounding space, as shown in the cut-open side views. **(F)** Workflow for HPF using the Autogrid planchettes including i) planchette preparation, ii) sample loading and vitrification and iii) grid retrieval under cryogenic conditions after freezing.

Complementary lid designs expand the versatility of the system: the “Waffle” lid, with a central protrusion that seals directly on top of the grid mesh, confines samples in solution between the grid bars, thereby limiting the sample thickness to the grid dimensions **(Figure 1C, D, Figure S1A)**. “Biopsy” lids incorporate a central, variable spacer, resulting in a total of 60 to 240 µm above the grid, enabling freezing of thicker specimens such as organoids, tissue samples or entire multicellular organisms, while preventing compression of the sample between the planchettes **(Figure 1C, E, Figure S1B)**. The spacer is positioned only over the grid mesh, restricting the sample to the grid area, excluding the Autogrid ring. Spacer dimensions can be adjusted to match sample thickness, allowing flexible adaptations to different specimen types **(Figure S1B, C-E)**. Since the Autogrid planchettes are compatible with conventional 6 mm planchettes, a flat planchette can be used as a lid for an intermediate thickness (∼120 µm on top of the grid mesh) **(Figure S1D)**.

All planchettes were iteratively optimized to improve handling and vitrification performance and were produced by our in-house workshops (see **Document S1** for technical drawings). Initial prototypes were manufactured from aluminum (AW-6082, AlSi1MgMn), a material commonly used for HPF planchettes **(Figure S1F-H)**. However, we observed practical limitations with this material, including difficulties in grid retrieval after freezing, potentially due to different thermal expansion between aluminum and the Autogrid during freezing. In addition, vitrification of the buffer surrounding cells was often not satisfactory. To address these issues, we transitioned to planchettes made from copper **(Figure 1B, C, Figure S1C-E)**. Copper more closely matches the material composition of Autogrids, which are likely composed of copper alloy, reducing mismatches in thermal expansion properties. Furthermore, copper has a substantially higher thermal conductivity (∼400 W/mK vs. ∼170 W/mK for aluminum 6082), improving heat transfer during freezing. Together, these properties improved both handling and vitrification performance in subsequent experiments.

HPF using Autogrid planchettes follows a streamlined workflow consisting of three main steps: (i) preparing the planchettes, (ii) sample loading and vitrification, and (iii) post-freezing grid retrieval **(Figure 1F, Video S1)**. (i) Surface treatment of the planchettes with 0.5% cetyl palmitate significantly improved grid release, as previously reported ^27^ **(Figure 1Fi)**. In addition, applying 1-hexadecene to the Autogrid planchette before adding the grid was crucial, as it filled all cavities of the planchette and prevented air from becoming trapped during planchette assembly, which would otherwise lead to deformation of the planchette or grid **(Figure 1Fi)**. (ii) After planchette preparation, clipped EM grids were loaded with sample, placed into the Autogrid planchette, and enclosed with the appropriate lid prior to vitrification using standard 6 mm HPF half-plates **(Figure 1Fii)**. (iii) Following vitrification, the planchettes were disassembled under liquid nitrogen, and the grid was retrieved using pointed tweezers via the side cutouts of the Autogrid planchette **(Figure 1Fiii)**. At cryogenic temperature, 1-hexadecene became brittle and fractured at the grid–planchette interface, facilitating efficient grid removal.

### Optimizing pFIB milling workflows for on-grid lamella preparation

To validate the freezing performance of the Autogrid planchette system, we froze *Saccharomyces cerevisiae* and the single-celled ciliate *Tetrahymena thermophila* using the Waffle lid configuration and transferred the samples to a plasma-FIB (pFIB) instrument equipped with an autoloader system, CompuStage and integrated light microscope. SEM overview images revealed homogenous ice thickness across the grid **(Figure 2A)**.

**Figure 2.**
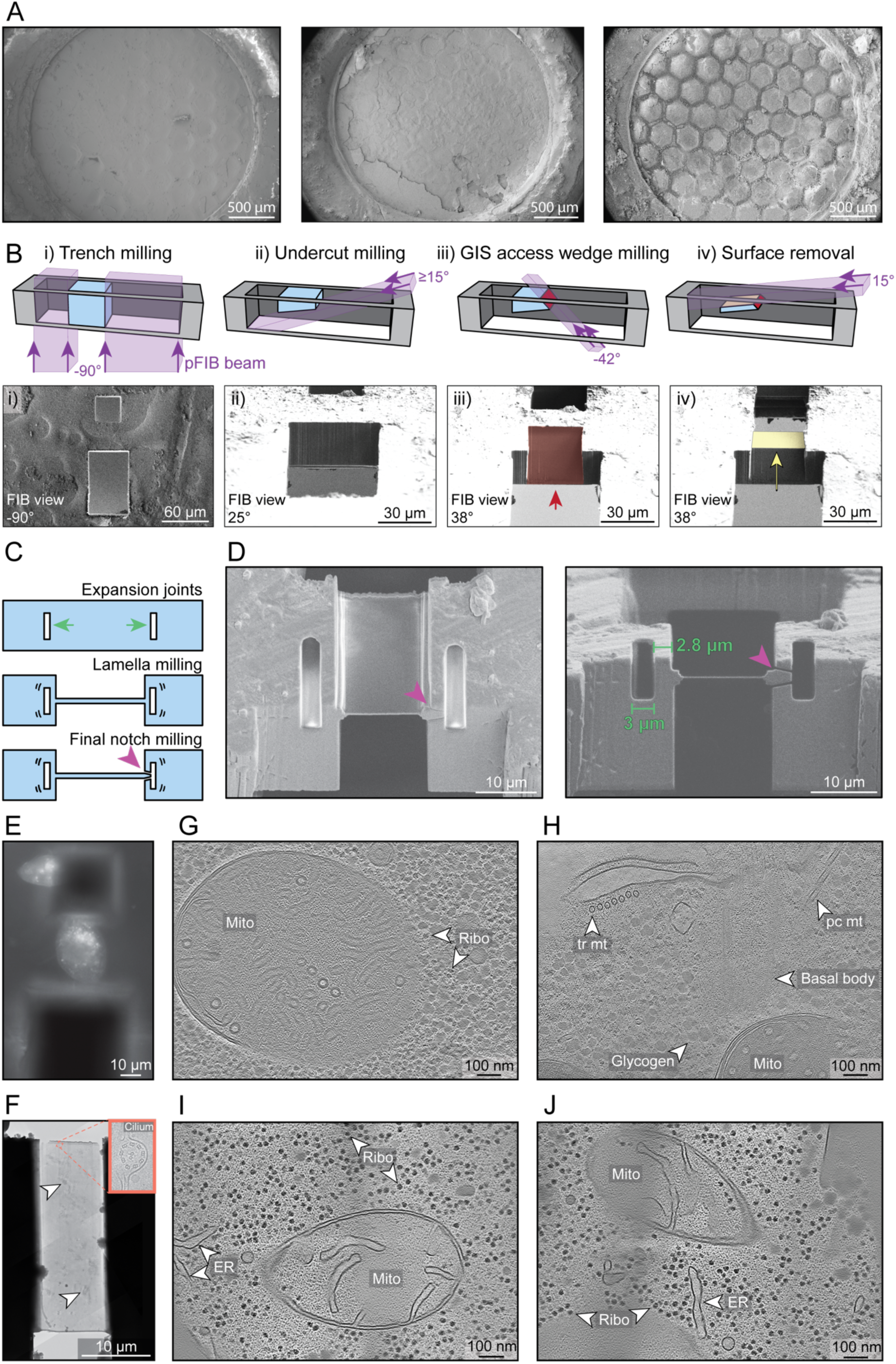
Optimizing pFIB milling workflows for on-grid lamella preparation. **(A)** SEM overview images of EM grids with *S. cerevisiae* (left panel) and *T. thermophila* (middle and right panel) cells, high-pressure frozen in Autogrid planchettes using the Waffle lid, showing homogenous ice thickness. **(B)** Schematic overview (top) and corresponding FIB images (bottom) of the on-grid lamella preparation workflow. Four preparation steps precede automatic lamella preparation: i) trench milling for access of the lamella preparation site, ii) undercut milling to remove material below the region of interest, iii) access wedge milling to generate an upward-facing surface for GIS deposition, and iv) surface removal to generate an even surface for GIS deposition and lamella milling. **(C)** Schematic of stress relief strategy during lamella preparation, combining expansion joints (green arrows) and detachment of the lamella from one side after final polishing (purple arrowhead). **(D)** Representative SEM (left) and FIB (right) images of an on-grid lamella prepared below the grid surface, using 3 µm wide expansion joints with an offset of 2.8 µm to the final lamella. The lamella was detached from one side after polishing (purple arrowhead). **(E)** Cryo-fluorescence light microscopy image of a high-pressure frozen *T. thermophila* cell after trench-milling. The auto-fluorescent signal was used to guide trench milling. **(F)** Cryo-TEM overview image of an on-grid lamella of *T. thermophila* prepared using expansion joints and lamella detachment for stress relief. Regions with crystalline ice reflections in the surrounding buffer are indicated by white arrowheads. The inset shows a zoom-in view to a cilium. **(G, H)** Slices through cryo-tomograms of *T. thermophila* cells demonstrate preserved ultrastructural features. The tomograms reveal the organization of organelles (Mito = Mitochondria) and macromolecular machines, such as a basal body and its associated transverse (tr mt) and post-ciliary microtubules (pc mt). Ribosomes (Ribo) and glycogen granules (Glycogen) are also clearly discernable. **(I, J)** Slices through cryo-tomograms of *S. cerevisiae* cells, showing preserved ultrastructure, including organelles such as mitochondria (Mito), endoplasmic reticulum (ER) and individual macromolecular complexes such as ribosomes (Ribo).

For on-grid lamella preparation, we optimized a Xenon-based pFIB-milling workflow. Surface ice contamination was first removed using a low-magnification, broad ion beam cleaning step, as previously described ^32^ **(Figure S2A)**. For auto-fluorescent samples such as *T. thermophila*, we identified regions of interest using the integrated fluorescence microscope to guide initial trench milling **(Figure 2E, Figure S2B)**. To allow for a large viewing angle, asymmetric trenches were used with a larger bottom trench (50 × 80 µm) and a smaller top trench (30 × 30 µm) **(Figure 2Bi, Figure S2C)**. Subsequent undercut milling was performed at gradually lower milling angles, until all material below the region of interest was removed **(Figure 2Bii)**.

As regions of interest were frequently located below the grid surface, we established a protocol for lamella preparation at varying depths. A homogenous layer of protective organometallic platinum using the gas injection system (GIS) is essential for minimizing curtaining and ensuring lamella survival ^38^. Conventional trench geometries, however, resulted in incomplete GIS coverage, particularly on the front edge in deeper regions, leading to characteristic triangular exclusion zones **(Figure S2D, left panel)**. To overcome this, a GIS access wedge was milled into the front face of the lamella at a −42° milling angle (milled from the back of the grid, **Figure 2Biii**), orienting the surface slightly more towards the GIS needle and enabling more homogenous organometallic platinum deposition on the leading edge **(Figure S2D, right panel)**. In addition, removal of the sample surface prior to GIS deposition generated a flatter deposition plane, reducing curtaining and enabling lamella preparation also in uneven samples **(Figure 2Biv, Figure S2E)**.

Another critical factor for lamella survival was stress relief. Due to the thickness of HPF samples, internal stresses frequently led to lamella breakage, a problem that may be amplified by mechanical coupling of the sample to the Autogrid. To mitigate this, we compared different stress-relief strategies, including expansion joints – commonly used for conventional lamella milling ^39^ – and notch milling, which was suggested recently for HPF samples ^28,31,32^. We tested different notch geometries, as well as expansion joints in combination with notch milling after the final polishing step ^32^ **(Figure 2C, Figure S2F)**. Expansion joints (3 µm wide) in close proximity (∼2.8 µm) to the final lamella, combined with detaching the final lamella from one side after polishing provided the most robust lamellae in our hands **(Figure 2D, Figure S3A)** ^40^. After lamella preparation, the grids were subsequently loaded into a cryo-transmission electron microscope **(Figure 2F)** for cryo-ET data acquisition.

All on-grid lamellae were prepared using samples frozen with the preliminary aluminum design of the planchettes, which yielded well-preserved cellular features in all specimens **(Figure 2G-J, Figure S3B, C)**. Intracellular organelles such as mitochondria, glycogen granules and the endoplasmic reticulum **(Figure 2G-J, Figure S3C)** as well as macromolecular machines such as ribosomes **(Figure 2I, J, Figure S3C)** and even higher order assemblies such as basal bodies and their associated microtubule bundles **(Figure 2H, Figure S3B)** were readily identified. Occasionally however, we observed crystalline ice in the surrounding buffer, indicating insufficient cryo-protection or incomplete vitrification **(Figure 2F)**. Together with limitations in throughput due to pFIB instabilities at our instrument, this motivated the transition to copper planchettes and the adoption of lift-out strategies for further evaluation of vitrification performance.

### Assessing vitrification performance of Autogrid planchettes by lift-out cryo-ET and subtomogram averaging

To benchmark the freezing performance of copper Autogrid planchettes, we froze *S. cerevisiae* cells using the Waffle configuration. The frozen grids were transferred to a pFIB-SEM instrument equipped with a cryo-micromanipulator and regions of interest were lifted out onto an acceptor grid using the SOLIST method ^33^ **(Figure 3A, Figure S4A-E)**. Multiple sample slices were obtained from every excised sample block **(Figure 3B)** and used for lamella preparation **(Figure 3C)**. As these slices were typically ∼5 µm thick and largely free of surrounding bulk material, lamellae could be prepared using established FIB-milling protocols without the need for additional stress relief adaptations. Typically, 5–20 lamellae could be generated within a single session **(Figure 3D)**. Once transferred to a cryo-TEM, individual *S. cerevisiae* cells were readily identified on lamella overview images **(Figure 3E)** and resulting cryo-tomograms revealed well-preserved cellular architecture, including mitochondria, vacuoles, the cell wall and plasma membrane, the endoplasmic reticulum and cytoplasmic complexes such as ribosomes **(Figure 3F-H, Figure S4F)**.

**Figure 3.**
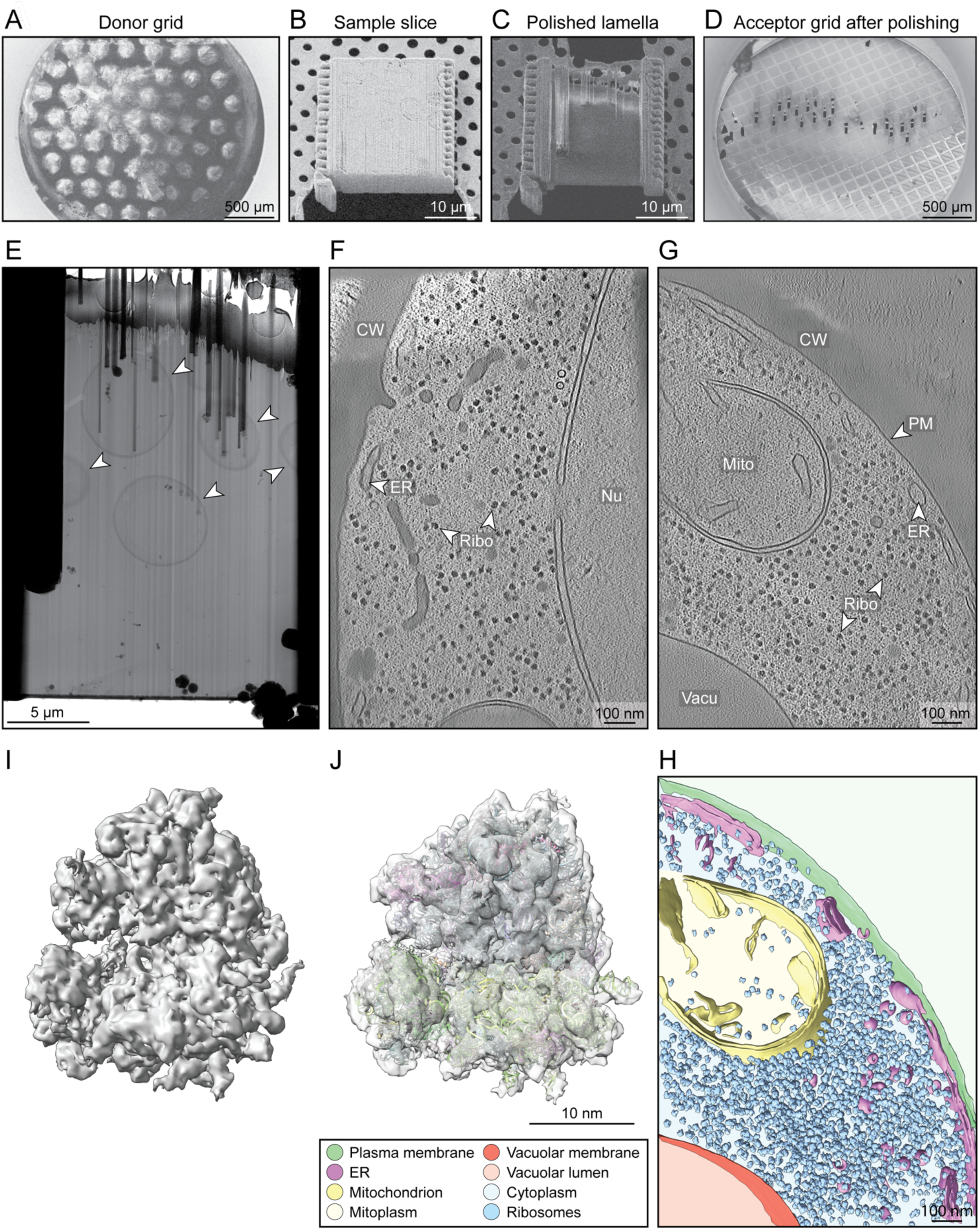
Assessing vitrification performance of Autogrid planchettes by lift-out cryo-ET and subtomogram averaging. **(A)** SEM image of an EM grid containing *S. cerevisiae* cells high-pressure frozen in 10% dextran using the Autogrid planchette system with a Waffle lid and subsequently used for cryo-lift-out. **(B)** SEM image of a sample slice lifted out onto an acceptor grid and attached to the grid by redeposition milling. The front side was polished to obtain a clean surface for sputter coating and GIS coating. **(C)** SEM image of the final lamella derived from the sample slice shown in (B). **(D)** SEM overview of the acceptor grid after lamella preparation. **(E)** Cryo-TEM overview image of the lamella shown in (C). Individual *S. cerevisiae* cells are indicated by white arrowheads. *S. cerevisiae* cells were vitrified in Autogrid planchettes using 10% dextran in YPD medium as cryo-protectant. No signs of incomplete vitrification were observed. **(F-G)** Slices through representative cryo-tomograms of *S. cerevisiae* cells showing well-preserved cellular ultrastructure. Organelles and protein complexes, such as ribosomes (Ribo) mitochondria (Mito), endoplasmic reticulum (ER) and vacuoles (Vacu), as well as the cell wall (CW) and plasma membrane (PM) are clearly resolved. Cells were vitrified in 20% BSA in PBS (F) or without any cryo-protectant in YPD medium (G) by HPF in Autogrid planchettes. **(H)** Three-dimensional segmentation of tomogram shown in (G). Cellular features are colored as indicated. **(I)** Subtomogram average of 80S ribosomes determined from *S. cerevisiae* tomograms, resolved to 8.8 Å resolution. **(J)** Structure of the 80S ribosomes fitted with the model of a yeast 80S ribosome (PDB: 8UT0).

To assess the vitrification capacities of our Autogrid planchettes, we analyzed lamellae and individual tilt-series for vitrification artifacts, such as Bragg reflections or ultrastructural damage caused by crystalline ice. As described above, the use of aluminum planchettes or sub-optimal cryo-protectants resulted in incomplete vitrification, particularly evident as Bragg reflections in the surrounding buffer, while overall cellular ultrastructure remained largely preserved across different sample types **(Figure 2F, Figure S5A)**. In contrast, switching to copper planchettes combined with the addition of 10% dextran as cryo-protectant substantially improved ice quality **(Figure 3E)**. Cryo-tomograms from copper planchette preparations revealed no signs of perturbed cellular architecture, with intact membranes without any evidence of membrane ruffling, consistent with a near-native state **(Figure 3F, G, Figure S4F)**. Fourier transform analysis of the corresponding tilt-series **(Videos S2-5)** revealed power spectra lacking reflections characteristic of crystalline ice **(Figure S5B)**, further supporting successful vitrification.

To further evaluate structural preservation beyond visual inspection, we performed subtomogram averaging of 80S ribosomes from 20 *S. cerevisiae* tomograms acquired from samples frozen in Autogrid planchettes. The averaged structure was resolved to 8.8 Å resolution **(Figure 3I, Figure S5C)** and aligned well with reference yeast ribosome models **(Figure 3J)**, indicating preservation of macromolecular integrity.

### High-pressure freezing of mouse brain biopsies in Autogrid planchettes conserves macromolecular features

As a proof-of-concept, we evaluated the feasibility of applying the Autogrid planchettes in the Biopsy configuration to tissue samples. Therefore, tissue fragments of mouse brain were obtained by manual dissection, transferred onto clipped Autogrids, and subjected to HPF using the Autogrid planchette in combination with a biopsy lid. The grids were subsequently transferred to a pFIB instrument with a micromanipulator **(Figure S6A)**, and lamellae were prepared using the SOLIST lift-out workflow as described above **(Figure S6B-E)**. We observed canonical features of neuronal cells in cryo-electron tomograms of pFIB-milled mouse brain biopsy material **(Figure S6F)**. While overall tissue integrity was partially compromised, likely due to extended pre-freezing handling times, subcellular features such as actin filaments **(Figure 4A)**, mitochondria **(Figure 4B, Figure S6F)**, presynaptic vesicles **(Figure S6F)** and filled microtubules **(Figure 4C)** were readily identifiable. Individual tubulin protofilaments were discernable within individual tomograms and in a subtomogram average, demonstrating structural preservation at a macromolecular level **(Figure 4D)**.

**Figure 4.**
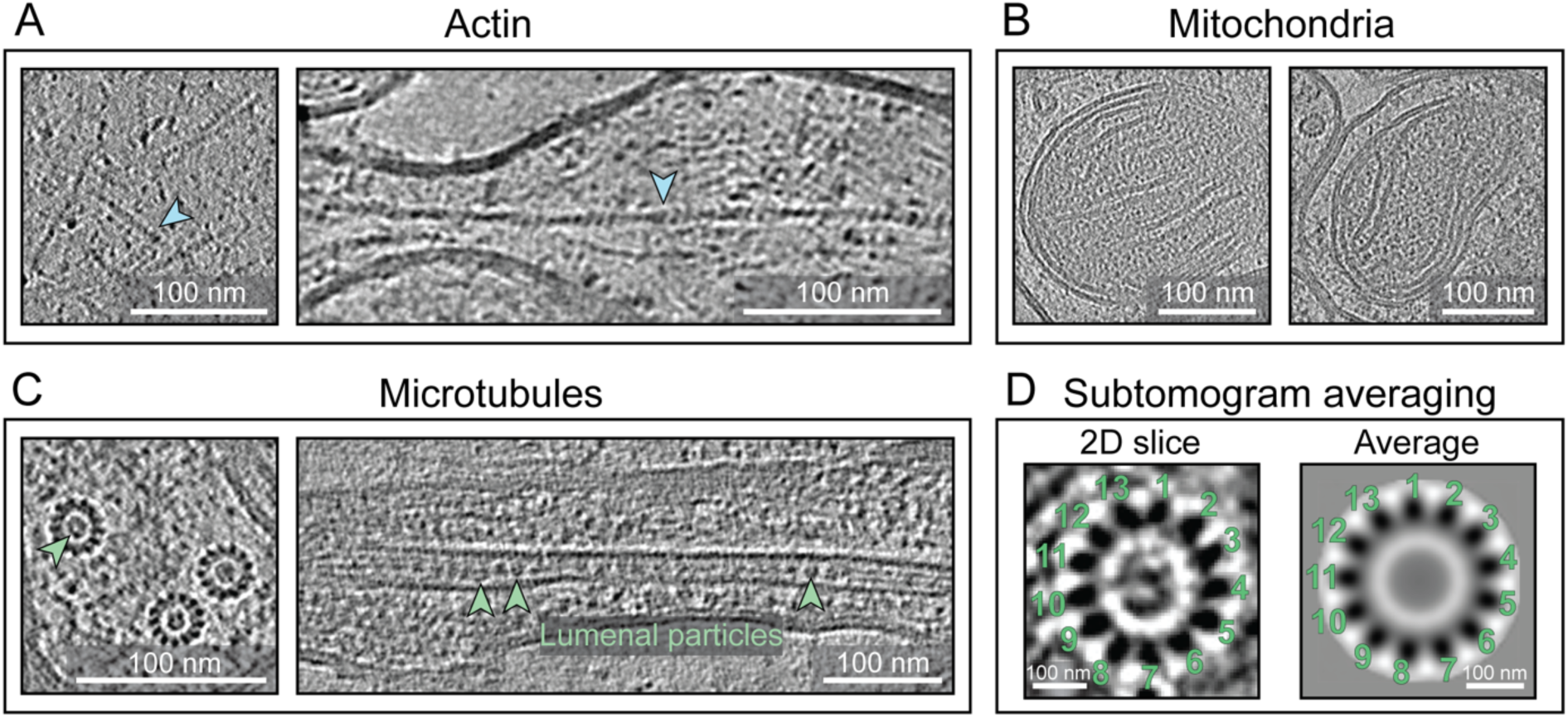
High-pressure freezing of mouse brain biopsies in Autogrid planchettes conserves macromolecular features. (A-C) Slices through cryo-tomograms of mouse brain biopsies frozen in Autogrid planchettes with the Biopsy lid configuration show preserved cellular structures, including actin filaments (A, blue arrowheads), mitochondria (B) and microtubules (C). Lumenal particles (green arrowheads) are visible within the microtubule lumen. **(D)** Cross-section of a microtubule within a cryo-tomogram of mouse brain tissue showing 13 protofilaments and luminal density (left). Subtomogram averaging confirms the protofilament architecture (right).

Together, these results demonstrate the feasibility of using the Autogrid planchettes for high-pressure freezing and subsequent downstream processing of tissue biopsies, supporting their potential for structural investigations of cellular architecture and macromolecular assemblies in complex multicellular specimens.

## Discussion

HPF, in combination with cryo-FIB milling and cryo-ET, enables structural investigation of complex biological systems beyond the thickness limit of plunge freezing. However, its broader adoption has been limited by workflow complexity and incompatibility with modern grid-based cryo-EM pipelines. Here, we address these challenges by introducing Autogrid planchettes that enable direct freezing of clipped EM grids, thereby simplifying handling and improving integration with downstream cryo-EM workflows. The preservation of cellular ultrastructure and ribosome structures at sub-nanometer resolution demonstrates that this approach maintains macromolecular integrity and is suitable for structural analysis by cryo-ET.

A key advantage of this system is the elimination of post-freezing grid handling steps, such as transfer and clipping, together with the improved mechanical stability of Autogrids, which collectively reduce grid damage and sample loss, while improving compatibility with automated cryo-FIB and TEM workflows. In addition, the modular design, combining an Autogrid bottom planchette with interchangeable lids, supports a wide range of sample types – from cellular suspensions to tissue specimens – and allows individual adaptation to specific experimental needs. Our technical drawings can serve as a framework to independently develop and customize the planchettes for specific requirements.

Our results highlight the importance of both device design and workflow optimization for reliable vitrification. Transitioning from aluminum to copper planchettes improved heat transfer and vitrification performance, while the use of appropriate cryo-protectants mitigated incomplete vitrification of the surrounding buffer. Recent systematic analyses on cryo-protectant performance provide a valuable resource for optimizing vitrification conditions across different sample types ^8,10^. Further improvements in planchette design may enhance performance even more, for example though the addition of a thin inert gold coating to minimize potential copper-related toxic effects on the sample and to improve surface properties. Despite these advances, certain limitations remain, including a potential mechanical coupling of samples to Autogrids, which may require adapted stress-relief strategies during on-grid lamella preparation.

Overall, the Autogrid planchette system provides a practical and scalable solution for integrating HPF into modern cryo-EM workflows. By simplifying sample preparation and improving robustness, this approach lowers the barrier for new users and expands access to structurally complex specimens, thereby supporting future applications of cryo-ET. In particular, for biomedical and potentially clinical research, reliable and reproducible workflows will be essential to unlock future diagnostic applications of cryo-ET.

## Materials and Methods

### Design and manufacturing of HPF planchettes

HPF planchettes were designed based on measurements of EM-grids clipped into commercial FIB-Autogrid rings using a scanning electron microscope. Planchettes were initially manufactured from Aluminum AW-6082 (AlSi1MgMn) by the physics workshop at ETH Zürich. After several rounds of iterations to improve dimensions and design, we switched to producing the planchettes from pure copper by the physics workshop at the University of Zürich. Biopsy lids were produced with different spacer cavities between of 240 µm, 200 µm, 120 µm or 60 µm above the grid, requiring a cavity of 120 µm, 80 µm, a flat surface, or a protrusion of 60 µm, respectively. Waffle lids were designed such that no spacer is present on top of the grid, requiring a central protrusion of 120 µm. We always calculated with a grid thickness of ∼20 µm.

STEP files of the optimized Autogrid planchette, the Waffle and the biopsy lids, as well as technical drawings can be found in the Supporting information.

### Cultivation of Saccharomyces cerevisiae

*Saccharomyces cerevisiae* were grown shaking at 25 °C and 180 rpm in YPD medium (10 g/l yeast extract, 20 g/l protease peptone, 20 g/l glucose) at pH 6.5 overnight. On the next day, the overnight culture was diluted to OD_600_ 0.1 – 0.2 AU in fresh YPD medium and grown until OD_600_ 1 AU was reached. Next, the culture was centrifuged at 800 x g for 3 min and the supernatant was removed. The pellet was resuspended in dextran or BSA cryo-protectants to a final concentration of 10-20% right before high-pressure freezing and centrifuged again at 800 x g for 3 min to generate a thick paste. The supernatant was removed and 3 µl of the pellet paste were added to clipped EM-grids as described below.

### Cultivation of Tetrahymena thermophila

*Tetrahymena thermophila* cells were grown shaking at room temperature in Modified Neff Medium (0.25% (w/v) protease peptone, 0.25% (w/v) yeast extract, 0.5% (w/v) glucose, 33.3 µM FeCl3). The cells at stationary phase were centrifuged at 300 x g for 5 min, the supernatant was removed, and the pellet was resuspended in 10 mM HEPES buffer at pH 7.5. Right before high-pressure freezing, the cells were mixed with dextran cryo-protectant to a final concentration of 10% and centrifuged again at 300 x g for 5 min. The supernatant was removed and several µl of the pellet were added to clipped EM-grids as described below.

### High-Pressure Freezing of Waffle samples in Autogrid planchettes

High-pressure freezing was performed on a Bal-Tec HPM100 (Bal-Tec AG) or Leica EM-ICE (Leica Microsystems) machine. All planchettes were coated with a solution of 0.5% cetyl palmitate in Chloroform by dipping the planchettes briefly into the solution and placing them on a cover slide to dry for ∼30 min. A drop of 1-hexadecene was added to the Autogrid planchettes right before freezing. EM grids (PELCO, hexagonal copper, 90 mesh) were clipped into FIB-Autogrid rings (Thermo Fisher Scientific) at room temperature and the clipped grids were glow discharged before use. 3 µl of yeast cells in cryo-protectant were applied to the grid. For Tetrahymena cells, 3 µl were applied and then blotted away with filter paper from below the grid. This procedure was repeated three times to increase cell number, while in the last round the cell solution was not blotted away. The clipped grid containing the sample was placed in the 1-hexadecene soaked Autogrid planchette and the sandwich was completed by addition of the Waffle lid planchette, with its protrusion facing the Autogrid planchette. The sample was immediately subjected to high-pressure freezing. Grids were retrieved from the planchettes by first removing the lid planchette and then retrieving the grid from the Autogrid planchette via the side cutouts using a tweezer. All handling steps were performed under liquid nitrogen and grids were stored in grid boxes until use.

### High-Pressure Freezing of mouse brain biopsies in Autogrid planchettes

High-pressure freezing was performed on a Bal-Tec HPM100 (Bal-Tec AG) machine. All planchettes were coated with a solution of 0.5% cetyl palmitate in chloroform by dipping the planchettes briefly into the solution and placing them on a cover slide to dry for ∼30 min. A drop of 1-hexadecene was added to the Autogrid planchettes right before freezing.

The brain of a C57BL/6 adult germ-free mouse was provided in fresh PBS buffer after extraction at ETH Zürich mouse facility (EPIC), Switzerland, as a byproduct of other experiments. All animal experiments were conducted in accordance with the institutional guidelines and Swiss federal regulations and were approved by the animal ethics committee of the Cantonal Veterinary Office, Zürich, Switzerland (permission no. ZH009/2021). For high-pressure freezing, the brain was dissected manually into slices and then trimmed into cubes of ∼200 µm with a scalpel. The biopsies were deposited onto EM grids (PELCO, hexagonal copper, 90 mesh) previously clipped in FIB-Autogrid rings (Thermo Fisher Scientific). The clipped grids containing the biopsies were placed in 1-hexadecene soaked Autogrid planchettes and the sandwich was completed by the addition of a Biopsy lid planchette with a 120 µm spacer, leading to a total volume of 240 µm above the grid. The sample was immediately subjected to high-pressure freezing and treated as described above.

### On-grid lamella preparation by plasma-focused ion beam milling

An Arctis plasma-FIB-SEM instrument (Thermo Fisher Scientific) was used for on-grid lamella preparation using xTUI and WebUI software. Ice contaminations were removed from the front and back side of the grid by FIB imaging at 15 nA at 30 kV for a few seconds on each position at the lowest possible magnification. Afterwards, the grid was sputter coated for 2 min at 70 nA, 12 kV current using the internal microsputter unit of the Arctis and covered with a protective layer of tri-methyl(methylcyclopentadienyl)platinum(IV) using a gas injection system (GIS) for 1 min. After GIS coating, a second layer of platinum was applied with the microsputter unit as described above. Regions containing cells of interest were identified using the integrated fluorescence light microscope (iFLM) module at a stage rotation of −180°. After regions of interest were identified, trench milling was performed from the backside of the grid at a stage angle of −128°, corresponding to a FIB angle of 90°. Two access trenches were milled above and below the regions of interest, leaving a region of ∼30 µm x 30 µm between the trenches. The bottom trench with a size of 50 µm x 80 µm and the top trench with a size of 30 µm x 30 µm were milled as rectangular patterns at 60 nA, 30 kV until all material was ablated. Afterwards, material below the lamella site was removed at gradually lower milling angles of 35°, 25°, 20° and 15° (corresponding to stage tilts of −3°, −13°, −18° and −23°, respectively) by milling rectangular patterns at 4 nA, 30 kV. Milling patterns covered the bottom trench region and the underside of the lamella, leaving the site for lamella placement untouched. Next, a wedge was milled into the front face of the lamella from the backside of the grid using a rectangular milling pattern at a milling angle of −42° (−80° stage angle) at 4 nA and 1 nA, 30 kV to generate an upwards facing slope for uniform GIS deposition. When the region of interest was located below the surface of the grid, the surface was removed and smoothed at a milling angle of 15° (−23° stage tilt) by milling a rectangular pattern at 4 nA and 1 nA, 30 kV. After all lamella sites were prepared, the grid was sputter coated again for 120 s at 70 nA, 12 kV, followed by 120 s of GIS deposition and an additional 120 s of microsputter coating at 70 nA, 12 kV. Afterwards, lamella milling was performed automatically using WebUI (Thermo Fisher Scientific). Lamellae were prepared at target milling angles of 15° – 20° (−18° to −23° stage tilt, respectively), and thicknesses of 150 – 220 nm were targeted. Expansion joints of 3 µm width with an offset of 2.8 µm from the final lamella were used for stress relief. After polishing, the backside of the lamella was thinned further by manual polishing at an overtilt of 1°. Finally, lamellae were detached on one side by milling line patterns that reach into the expansion joints at 0.1 nA, 30 kV, disconnecting the lamella from the surrounding material on one side for further stress relief.

Alternative milling procedures were assessed as well: Stress relief notches of two different geometries were milled at 1 nA, 30 kV after lamella site preparation, but before automatic lamella milling in WebUI. A rectangular notch geometry as well as an arrow-shaped notch geometry were assessed; the geometry can be seen in Figure S2Fi and ii.

### Lamella preparation by plasma-focused ion beam milling using SOLIST Lift-out

A Hydra Bio plasma-FIB-SEM instrument (Thermo Fisher Scientific) equipped with an EasyLift system was used for lamella preparation using lift-out with the SOLIST method, as described previously ^33^. Therefore, the EasyLift needle was modified by attaching a gold block adaptor using redeposition milling as previously described ^33^. The gold block was cut out from the UltrAuFoil (R2/2, 200 mesh, Quantifoil) acceptor grid. After gold block attachment, the donor grid was prepared. Ice contaminations were removed from the front side of the grid by FIB imaging at 15 nA at 30 kV for a few seconds on each position at the lowest possible magnification. Afterwards, the grid was sputter coated for 120s at 0.12 µA, 12 kV using the internal microsputter unit of the Hydra and covered with a protective layer of organo-metallic platinum using a gas injection system (GIS) for 90 s. After GIS coating, a second layer of platinum was applied with the microsputter unit as described above.

Afterwards, the sample volume that should be used for Lift-out was prepared as follows: with the sample perpendicular to the pFIB beam, a ∼50 µm wide and 80-120 μm long trench at the front and 10 μm wide trenches on all other sides were milled around the region of interest at 15-60 nA, 30 kV, leaving a 10 μm wide bridge on one side, connecting the block to the rest of the sample. The sides of the block were then polished at 1-4 nA, 30 kV. Undercut milling was performed on the bridge, starting at a milling angle of 50° and ending at a milling angle of 9°, using a current of 1 nA at 30 nV. The surface of the sample block was polished at 9° milling angle with 1 nA, 30 kV, to generate a smooth surface for needle attachment. Next, the EasyLift needle with gold-block attachment was inserted and lowered onto the surface at the position of interest. The gold block was attached to the sample block by redeposition milling, by placing cleaning cross section patterns on the gold block and milling from bottom to top, to re-deposit material into the cavity between the gold and the sample block, as previously described ^33^. Afterwards, the sample block was released from the bulk sample by milling away the remaining bridge connection at 1 nA, 30 kV. The sample block was lifted out of the donor grid and retracted with the EasyLift needle. The stage was moved to the acceptor grid. As acceptor grid, a 200 mesh UltrAUfoil grid (Quantifoil) was used, as described previously ^33^. Before insertion into the pFIB instrument, the acceptor grid was glow discharged for 120s at 20 mA. To attach the sample block to the acceptor grid, the stage was tilted to 6 – 11° milling angle for optimal attachment. The sample block was lowered onto a square of the UltrAUfoil grid until it touched down on the gold foil, then the needle was lifted slightly, and the lower ∼5 µm of the sample block were cut off using a line pattern at 1 nA, 30 kV. The remaining sample block was transferred to the neighboring square and the procedure was repeated. After all slices were prepared, they were attached to the gold foil by milling microstiches on the left and right sides of the slice at a milling angle of 90° as previously described ^33^. The front side of the slices was polished at 1 nA and 0.3 nA, 30 kV, microsputter coated with platinum for 120 s at 0.12 µA, 12 kV, and GIS coating was performed three times from different angles: 20s at 30° milling angle, 20 s at 90° milling angle, and again 20s at 30° milling angle. Afterwards, the grid was microsputter coated again as described above.

Finally, target positions were mapped using Maps 3.29 (Thermo Fisher Scientific) and lamellae were thinned to target thicknesses of 100 – 110 nm using AutoTEM Cryo 2.4.4 (Thermo Fisher Scientific), resulting in lamellae thicknesses of ∼200 nm.

### Cryo-ET data collection

The grids were analyzed using two 300 kV Titan Krios electron microscopes (Thermo Fisher Scientific) equipped with K3 direct electron detectors (Gatan) mounted on post-column energy filters (Gatan). Tilt series were acquired in zero-loss energy mode with a 20 eV slit at a magnification of 33’000 x, resulting in a pixel size of 2.678 Å/px or 2.699 Å/px depending on the microscope. SerialEM 4.2.0 and 4.2.0beta in low dose mode were used for data acquisition ^41^. PACEtomo 1.7.0 ^42^ and SPACEtomo 1.3 ^43^ were used for identifying target positions on the lamellae and collecting tilt series with a dose-symmetric tilt scheme ^44^ with a start tilt of −9°, a tilt range of −69° to 51° and an increment of 3°, corresponding to 41 projections per tilt series and a total accumulative electron dose of ∼120 e/Å^2^ at a defocus of −3 to −6 µm.

### Cryo-ET data processing

Tilt series were processed using the IMOD 5.1 software package ^45^. Frame alignment was performed using alignframes and tomograms were reconstructed using etomo with patch tracking. For visualization purposes, tomograms were filtered using DeepDeWedge ^46^ and IsoNet2 v2.0.0-beta ^47^. Cellular structures present in the tomograms were segmented using Dragonfly (Comet Technologies Canada; v. 2025.2) as previously described ^48^. Segmentation results were visualized using UCSF ChimeraX ^49^.

### Subtomogram averaging

Subtomogram averaging of ribosomes was performed on 20 cryo-tomograms of *S. cerevisiae* cells high-pressure frozen in Autogrid planchettes using the Waffle lid and prepared by on-grid lamella preparation. For subtomogram averaging, ribosome particles were processed using a Warp-RELION-M pipeline as previously described ^50^. In brief, tilt series were pre-processed using WarpTools, including motion correction, CTF estimation and handedness check. Next, tomograms were reconstructed with Warp and denoised with cryoCARE ^51^. Individual ribosome particles were picked using Cryolo ^52^, converted to RELION-3.0 star format and subtomograms were exported using WarpTools at a pixel size of 8 Å. Subtomograms were afterwards used for 3D classification in RELION 3.0 ^53^. Particles in 3D classes showing ribosome features were subjected to 3D refinement in RELION and then used for M refinement at a pixel size of 3 Å, including image warp grid, volume warp grid, particle poses and stage angles. A final resolution of 8.8 Å was obtained from 18’911 particles after post-processing in RELION 3.0 based on gold-standard FSC ^54^, which was applied with a b-factor of −50.

For subtomogram averaging of microtubules, individual microtubule filaments were manually picked in tomograms reconstructed with IMOD 5.1. For picking, the “filamentWithTorsion” model in Dynamo^55^ was used with an interbox distance of 21.4 nm. The coordinates and orientations were used to extract 2’489 individual microtubule particles as sub-tomograms at a binning factor of 4, corresponding to a pixel size of 10.71 Å/pixel, with a box size of 34.3 nm using the dtcrop function. The cropped particles were subjected to subtomogram averaging with Dynamo, using a random average of 50 particles as reference. The refined orientations were used to crop the particles again at a binning factor of 2 from CTF-corrected tomograms. The dataset was split in halves, and each dataset was refined individually, applying a C13 symmetry. The resolution of the resulting structure was determined by Fourier shell correlation using the dfsc function in Dynamo.

## Supporting information

Technical Drawings

3D Model files

## Resource availability

### Lead contact

Requests for further information and resources should be directed to and will be fulfilled by the lead contact, Gregor Weiss.

### Materials availability

Technical drawings and 3D model files of Autogrid planchettes are available in the supplementary material or upon request. Planchettes for test purposes are available from the lead contact depending on availability and may require contribution to production costs.

### Data and code availability

Representative cryo-ET data and ribosome subtomogram average have been deposited in the Electron Microscopy Data Bank as EMD-57483 to EMD-57502. This paper does not report original code. Any additional information required to reanalyze the data reported in this paper is available from the lead contact upon request. Figure 1 was partially created in BioRender. Weber, M. (2026) https://BioRender.com/eua2kir.

## Acknowledgements

The authors would like to thank the physics workshops at ETH Zürich and UZH Zürich for their valuable ideas about planchette design and the production of the planchettes. Martin Pilhofer is acknowledged for support throughout the project. The authors would also like to thank ScopeM at ETH Zürich and the Center for Microscopy and Image Analysis at the University of Zurich for providing support and equipment, especially Imre Gonda for sharing his expertise on the SOLIST method, Piotr Szwedziak for help with data acquisition and Andres Kaech for sharing his expertise on high-pressure freezing and planchette design. The authors would like to thank Bin Cai and Michal Wieczorek for providing *Tetrahymena thermophila* samples and Jannik Hugener for providing *Saccharomyces cerevisiae* samples. The authors would like to thank Selma Aslani, Yagmur Turgay and Emma Slack for providing mouse brain tissue. This research was supported by a Swiss National Science Foundation Starting Grant (TMSGI3_226208) and the Jean-Jacques et Letitia Lopez-Loreta Foundation.

## Author contributions

M.S.W and G.L.W designed the Autogrid planchette system. M.S.W. and T.E. prepared the samples, performed the HPF and pFIB experiments, recorded and analyzed data. J.X. performed subtomogram averaging of ribosomes. G.L.W. provided resources, funding and administration. M.S.W. together with G.L.W. conceived the research and wrote the manuscript.

## Declaration of generative AI and AI-assisted technologies in the writing process

During the preparation of this work the authors used ChatGPT 5 in order to improve readability and language of the manuscript. After using this tool/service, the authors reviewed and edited the content as needed and take full responsibility for the content of the published article.

---

**Figure S1.**
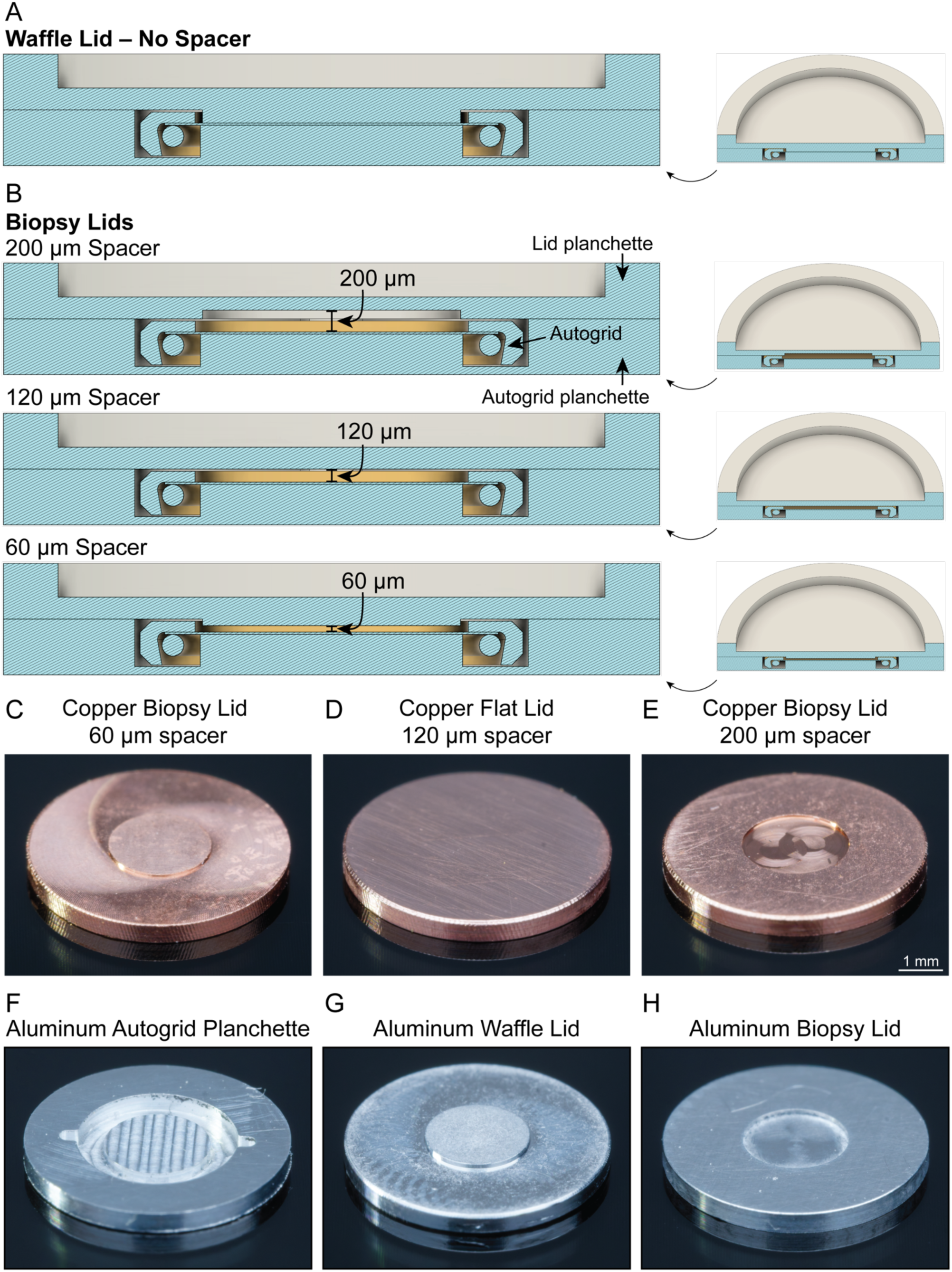
The Autogrid Planchette system and lid dimensions. **(A)** Schematic cut-open view of the Autogrid Planchette system with the waffle lid and a clipped EM-Autogrid. **(B)** Schematic cut-open views of the Autogrid Planchette system with a clipped EM-Autogrid and three biopsy lids with different spacers. **(C-E)** Example photographs of three different copper biopsy lid planchettes, designed to accommodate 60 µm (C), 120 µm (D) or 200 µm thick samples (E). **(F-H)** Example photographs of the aluminum Autogrid planchette (F), the Waffle lid (G), and a biopsy lid with a spacer of 240 µm (H).

**Figure S2.**
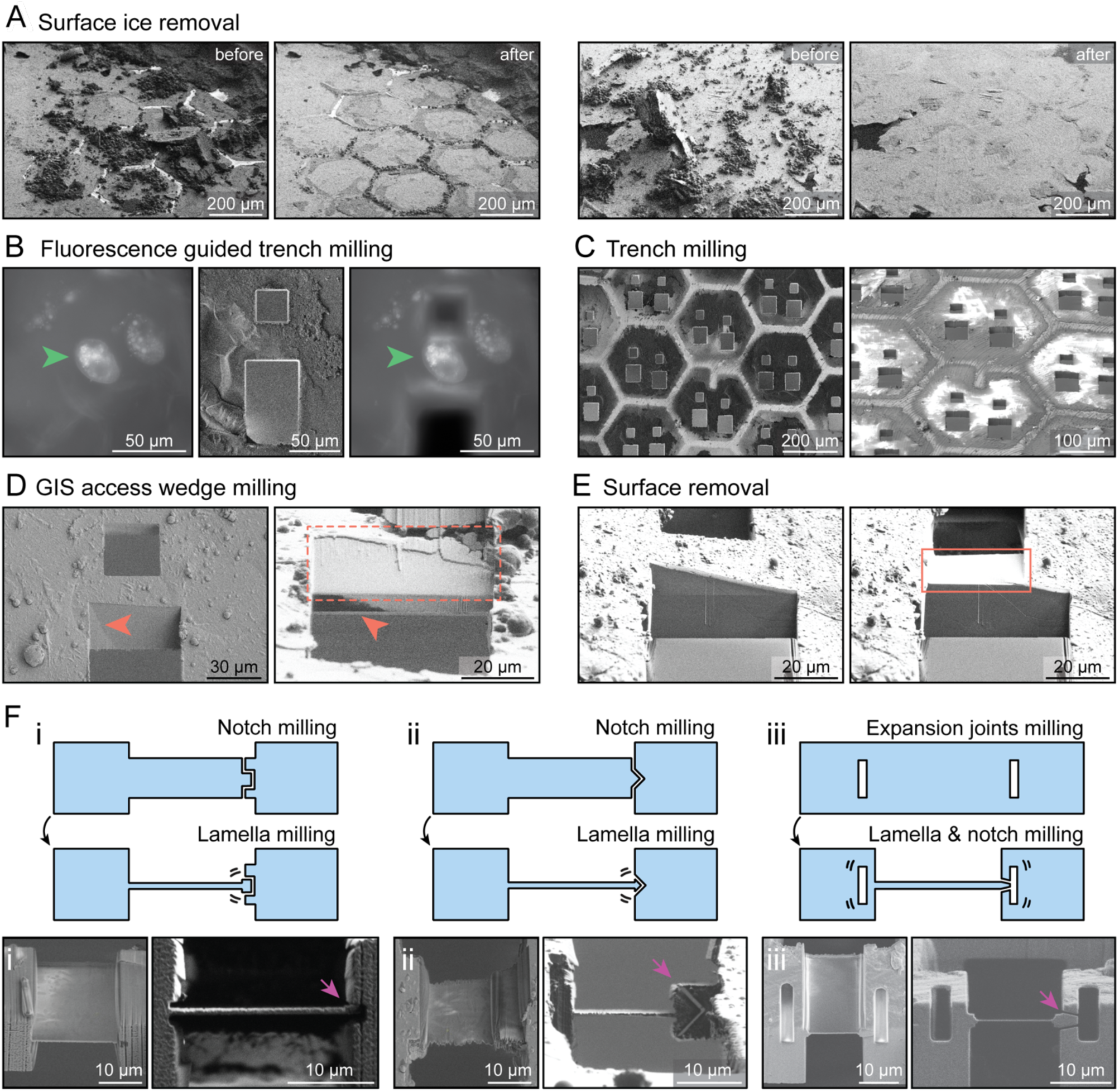
Technical details of the on-grid lamella preparation workflow. **(A)** SEM overview images before (left panels) and after (right panels) surface ice removal by a low-magnification broad ion beam. **(B)** Fluorescence guided trench milling of *T. thermophila* cells. Left and right panels show images taken with the integrated fluorescence microscope of the pFIB/SEM instrument. Middle panel shows FIB image post-trench milling. Green arrowheads point to the same cell before and after trench milling. **(C)** SEM images of EM grid at different angles post trench milling. Multiple trenches were milled to prepare lamella sites. **(D)** FIB/SEM images demonstrating uniform GIS deposition after access wedge milling. Left panel shows a SEM image with a triangular exclusion zone (orange arrowhead) after GIS deposition without an access wedge. Right panel shows a FIB image revealing uniform GIS deposition in the area opened with an access wedge (orange dotted box), while the region not included in the access wedge still shows an exclusion zone (orange arrowhead). **(E)** FIB images before and after the removal of the sample surface to enable uniform GIS deposition and generating a flat surface for lamella preparation (orange box). **(F)** Schematics (top), SEM images (bottom left) and FIB images (bottom right) of different notch milling strategies that were assessed for stress release. Rectangular (i) and arrow-shaped (ii) notches were milled before automatic lamella preparation. Preparing expansion joints before automatic lamella preparation, in combination with detaching the lamella from one side after final lamella milling (iii) resulted in the highest lamella survival rate. Purple arrowhs indicate point of stress release.

**Figure S3.**
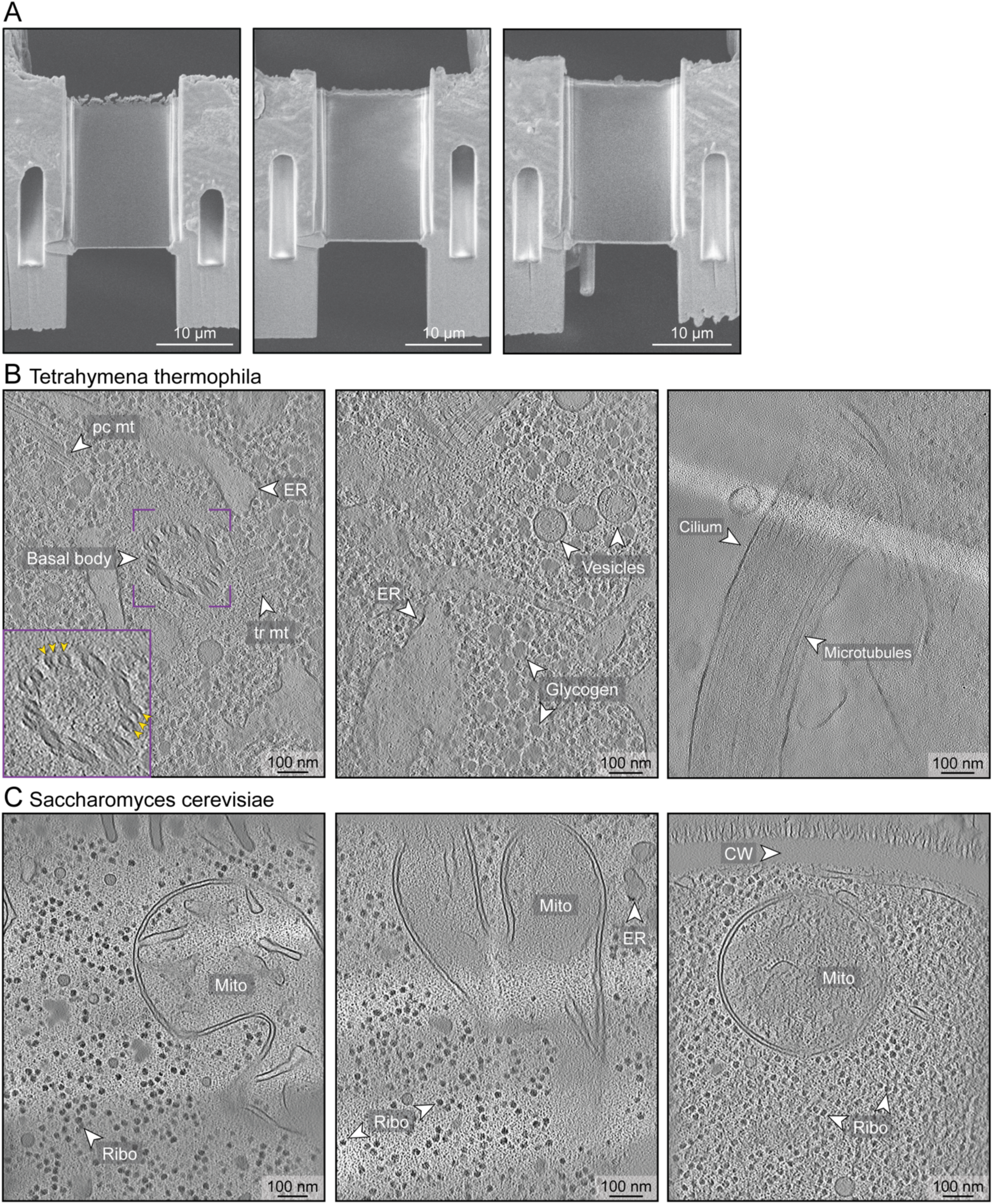
Example lamellae and cryo-tomograms prepared by on-grid lamella preparation. **(A)** Representative SEM images of *S. cerevisiae* lamellae prepared using expansions joints in combination with detaching one lamella edge after polishing. **(B)** Slices through cryo-tomograms of *T. thermophila* cells, revealing well-preserved cellular landscape including organelles, macromolecular machines and individual proteins. Purple box shows a zoom-in to a basal body, revealing individual microtubule triplets (orange arrowheads). Mito = mitochondria, Ribo = ribosomes, pc mt = post-ciliary microtubules, tr mt = transverse microtubules, ER = endoplasmic reticulum, Glycogen = glycogen granules. **(C)** Slices through cryo-tomograms of *S. cerevisiae* cells revealing a near-native preservation of cellular features. Mito = mitochondria, Ribo = ribosomes, ER = endoplasmic reticulum, CW = cell wall.

**Figure S4.**
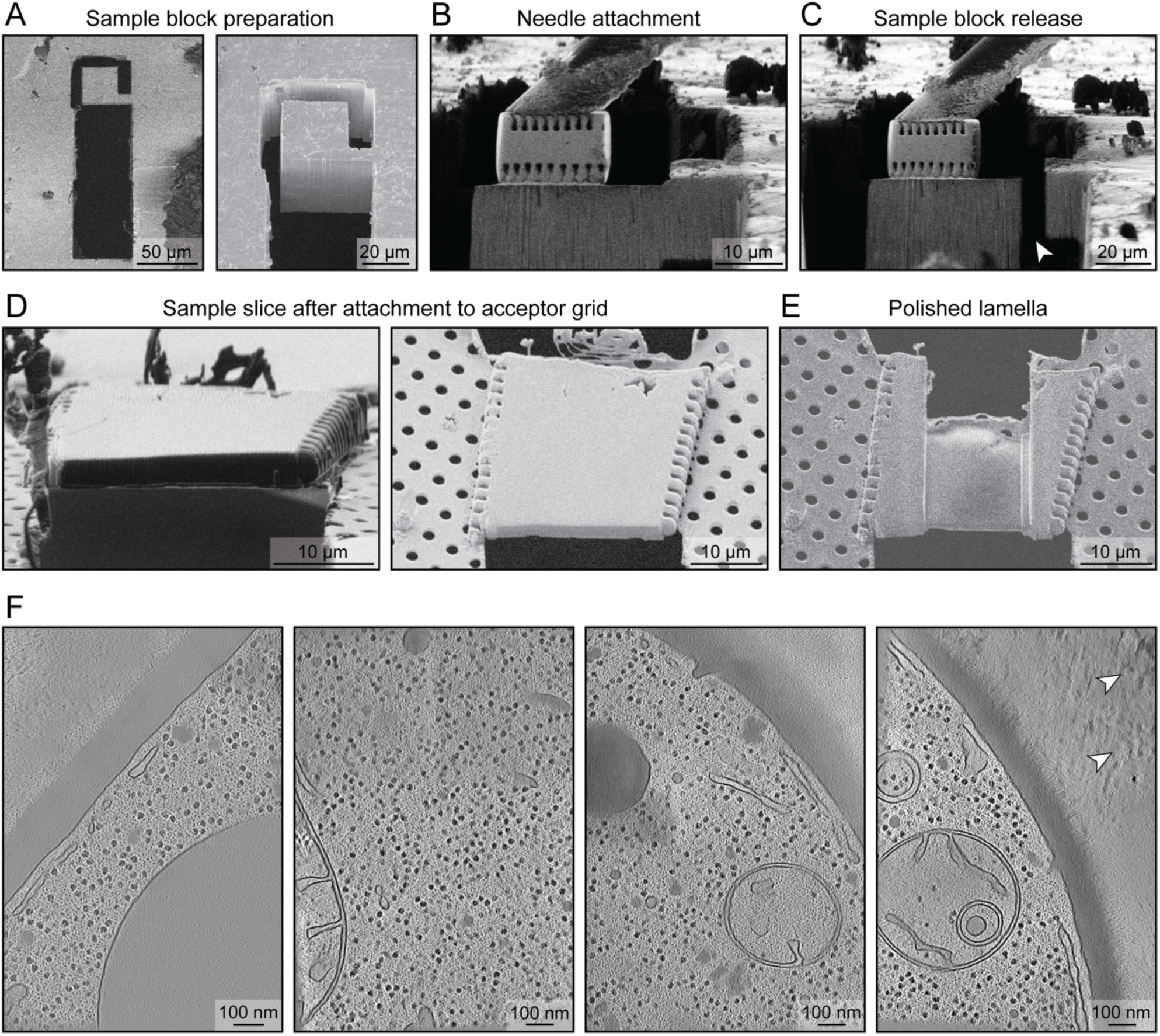
Cryo-lift-out workflow for samples frozen with the copper Autogrid planchette system. **(A)** SEM overview image of a sample block prepared for lift-out. **(B)** FIB image showing attachment of the micromanipulator needle to the sample block. A gold block adapter is used to connect the EasyLift needle to the specimen. **(C)** FIB image of the sample block after release from the surrounding bulk material. **(D)** FIB (left) and SEM (right) images of sample slice after attachment to the acceptor grid. **(E)** SEM image of final lamella from the sample slice shown in (D). **(F)** Representative cryo-tomograms of *S. cerevisiae* cells high-pressure frozen with the Autogrid planchette system in the Waffle lid configuration. Occasional crystalline ice reflections (white arrowheads) are visible in the surrounding buffer in one tomogram, likely due to suboptimal cryo-protectant conditions (20% BSA in PBS).

**Figure S5.**
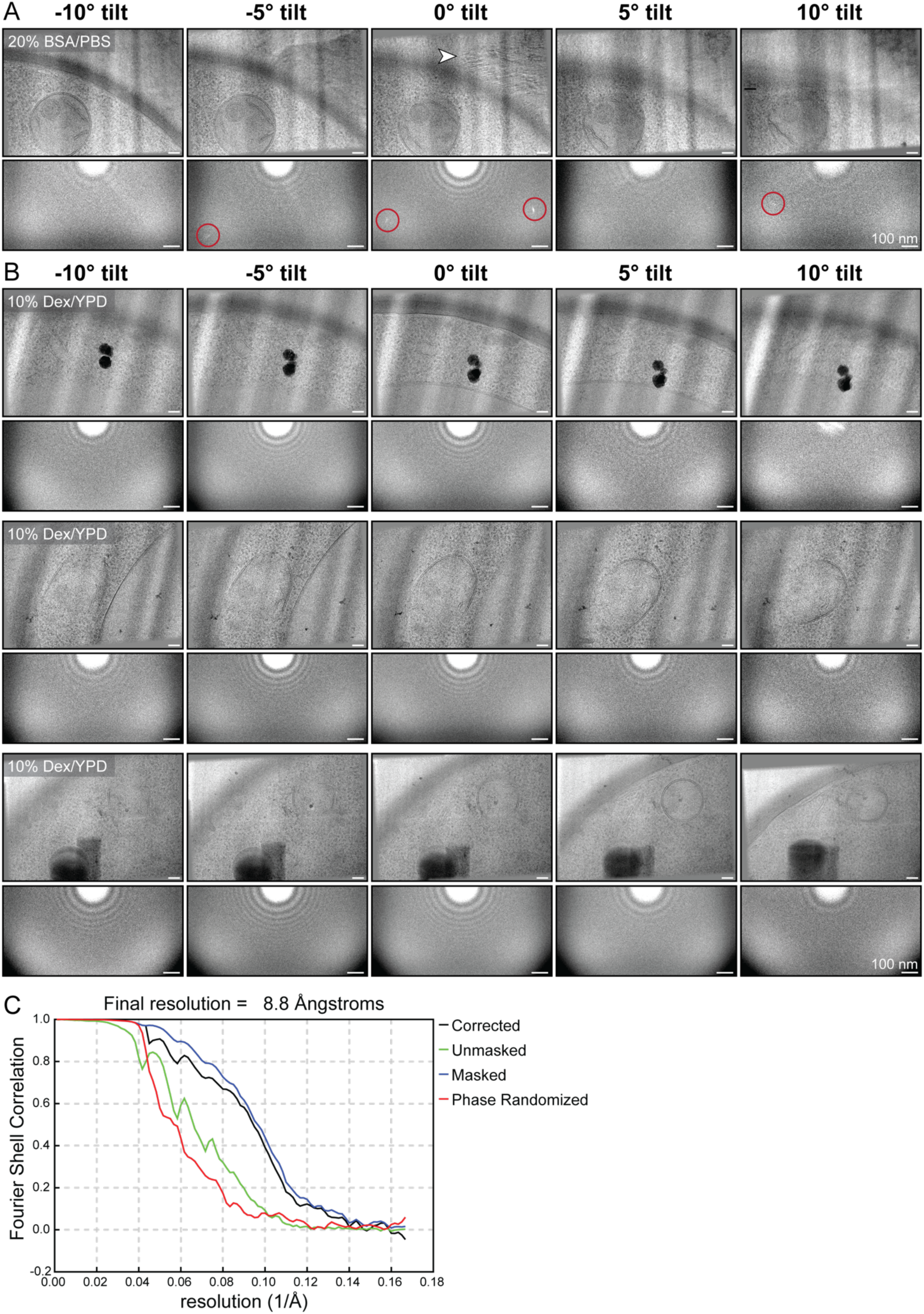
Analysis of vitrification quality in copper Autogrid planchettes. **(A)** Cryo-TEM projection images of an *S. cerevisiae* cell at different angles (top) and corresponding power spectra (bottom). Freezing *S. cerevisiae* in 20% BSA in PBS yielded incomplete vitrification, visible as Bragg reflections in the projection images and corresponding ice diffraction spots in the power spectra (red circles). Also see Video S2. **(B)** Cryo-TEM projection images of *S. cerevisiae* cells at different angles (top) and their corresponding power spectra (bottom). Freezing *S. cerevisiae* in 10% dextran in YPD medium resulted in effective vitrification, with no detectable Bragg reflections in projection images or diffraction spots in the power spectra. Also see Videos S3-5. **(C)** Gold-standard Fourier shell correlation (FSC) curve of the *S. cerevisiae* 80S ribosome subtomogram average. The map could be refined to a resolution of 8.8 Å.

**Figure S6.**
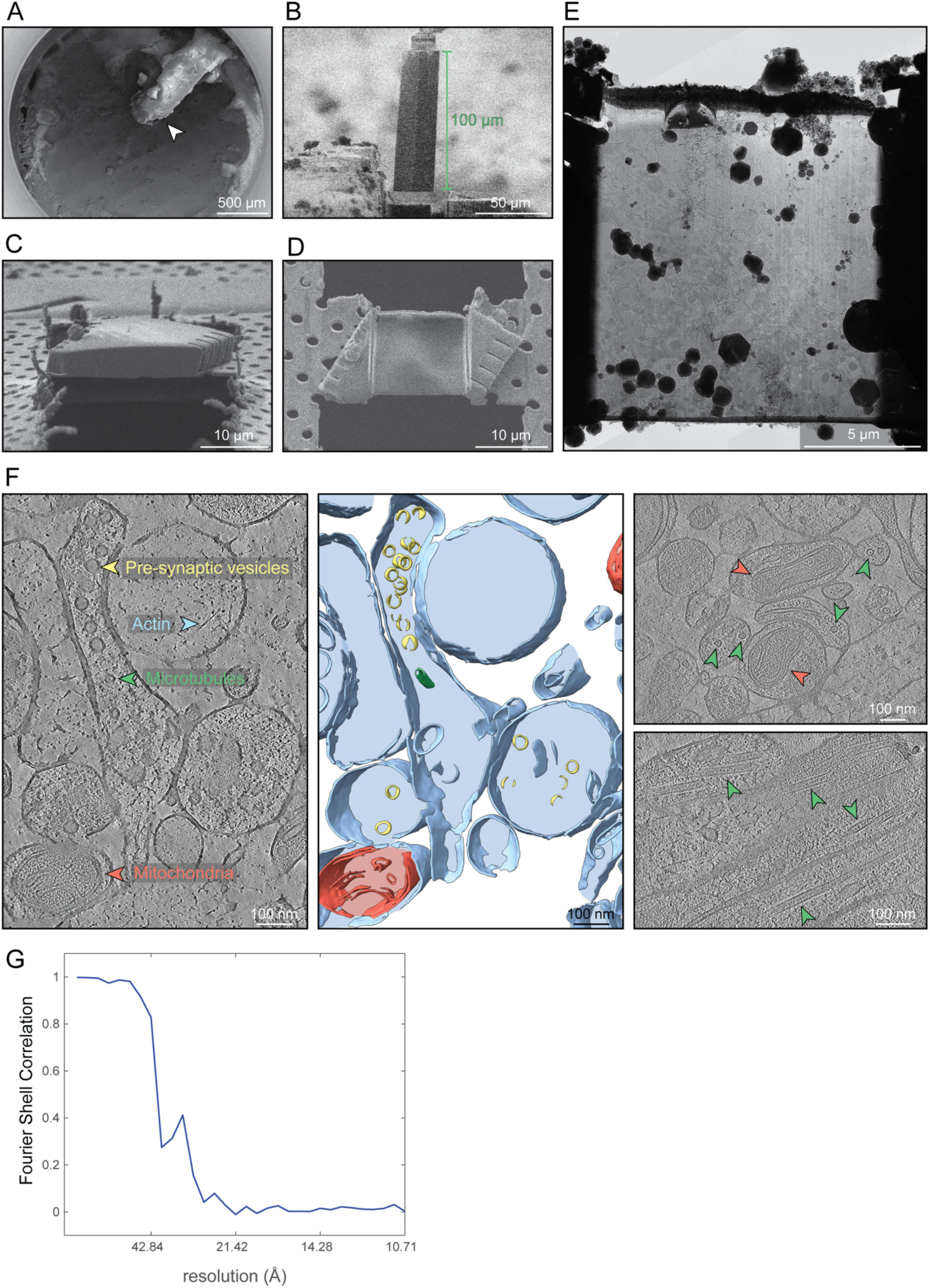
Cryo-ET of mouse brain biopsy frozen in the Autogrid planchettes. **(A)** SEM overview image of an EM-grid containing a mouse brain tissue (arrowhead), high-pressure frozen using the aluminum Autogrid planchette system in the Biopsy lid configuration. **(B)** FIB image of tissue block lifted out from the donor grid. **(C)** FIB image of a sample slice deposited on the acceptor grid. **(D)** SEM image of final lamella derived from sample slice shown in (C). **(E)** Cryo-TEM overview image of the lamella shown in (D). Cellular features can be readily identified on the lamella. **(F)** Representative cryo-tomograms and corresponding 3D segmentation of mouse brain tissue. Individual cells (blue) are visible, containing subcellular features such as pre-synaptic vesicles (yellow), actin filaments (blue arrowhead), mitochondria (red, red arrowheads) and microtubules (green, green arrowheads). **(G)** Fourier shell correlation (FSC) curve of the microtubule subtomogram average. The map could be refined to a resolution of ∼30 Å.

