## Supplementary material for "A simplified High-Pressure Freezing Workflow in Autogrids": Technical Drawings

### Autogrid Planchette

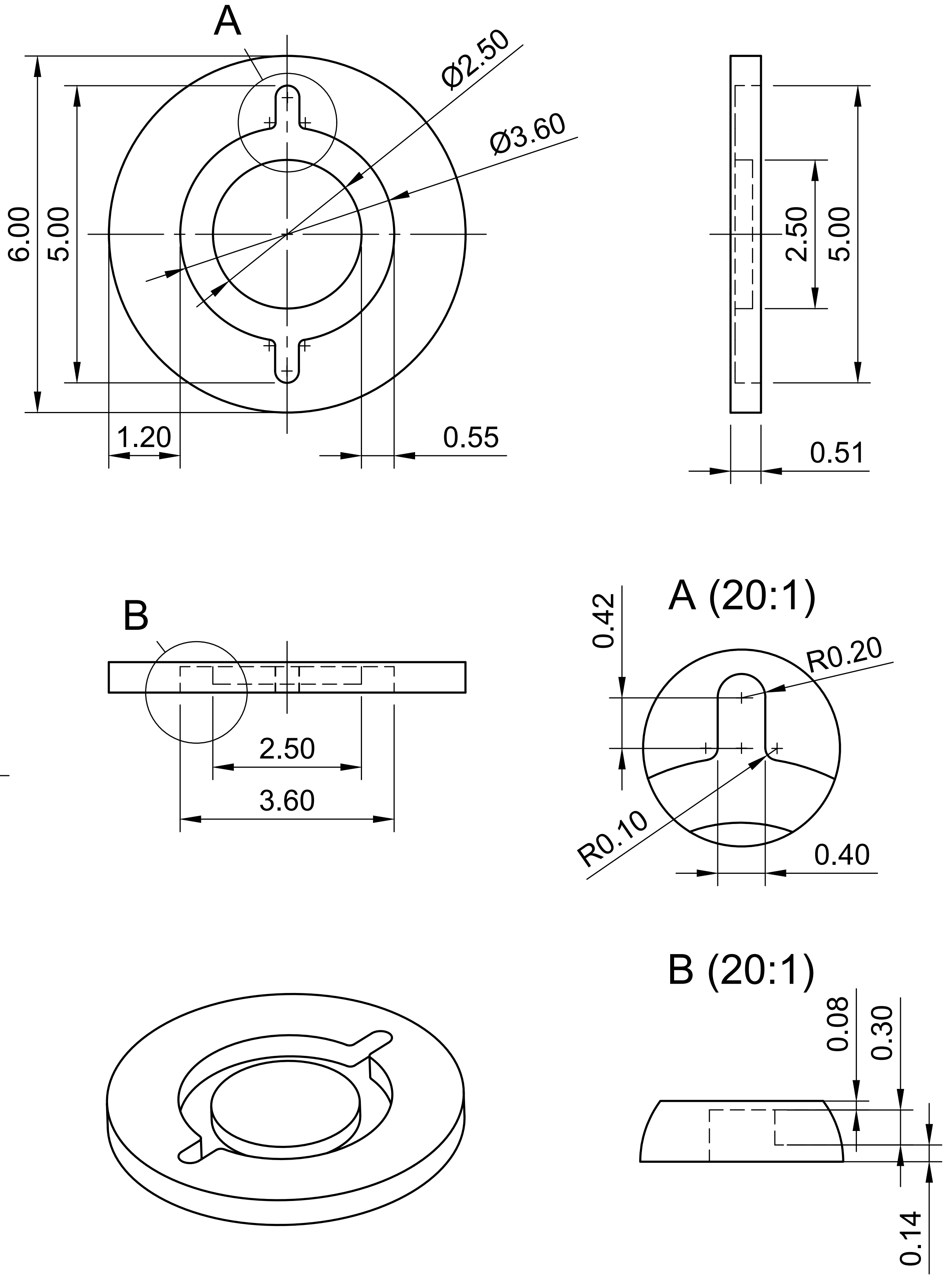

| Dept. | Technical reference | Created by | Approved by |
|---|---|---|---|
| | | Miriam Weber 14.04.2026 | |

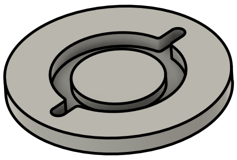

| | |
|---|---|
| Document type | Document status |
| Title: Autogrid_Planchette | DWG No. |
| | Rev. / Date of issue / Sheet: 1/1 |

### Waffle Lid - No spacer

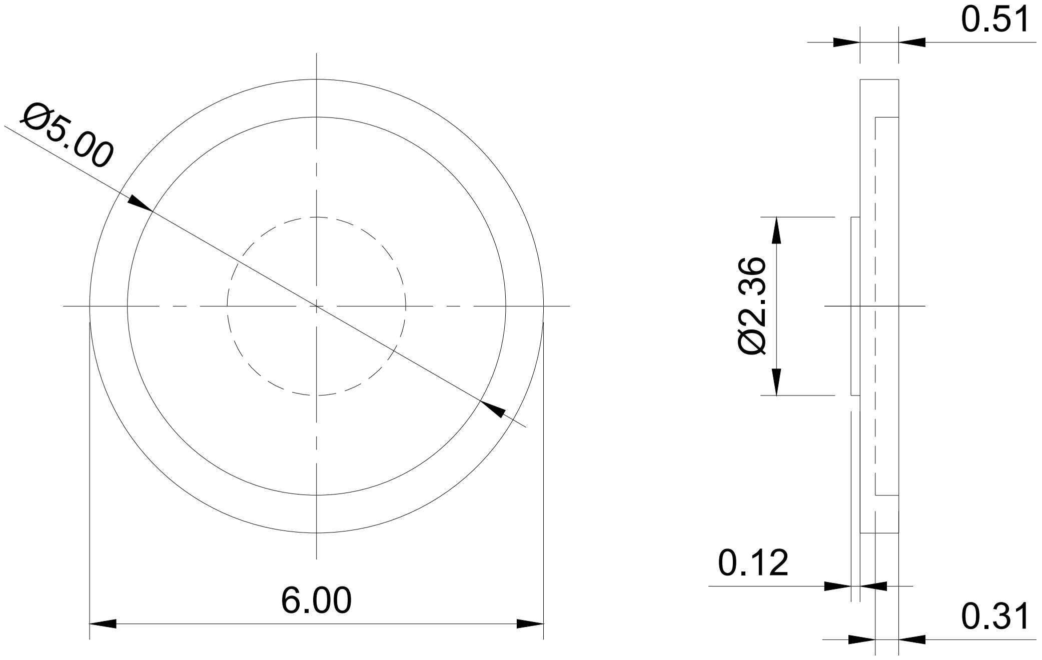

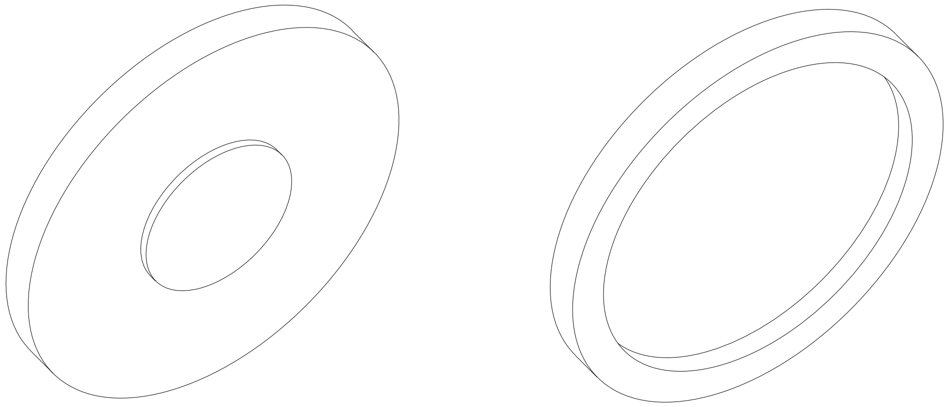

| Dept. | Technical reference | Created by | Approved by |
|---|---|---|---|
| | | Miriam Weber 14.04.2026 | |

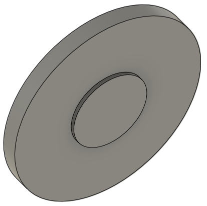

| Document type | Document status |
|---|---|

| Title | DWG No. | | |
|---|---|---|---|
| Waffle_Lid_no_spacer | | | |
| | Rev. | Date of issue | Sheet |
| | | | 1/1 |

### Biopsy Lid - 200 µm spacer

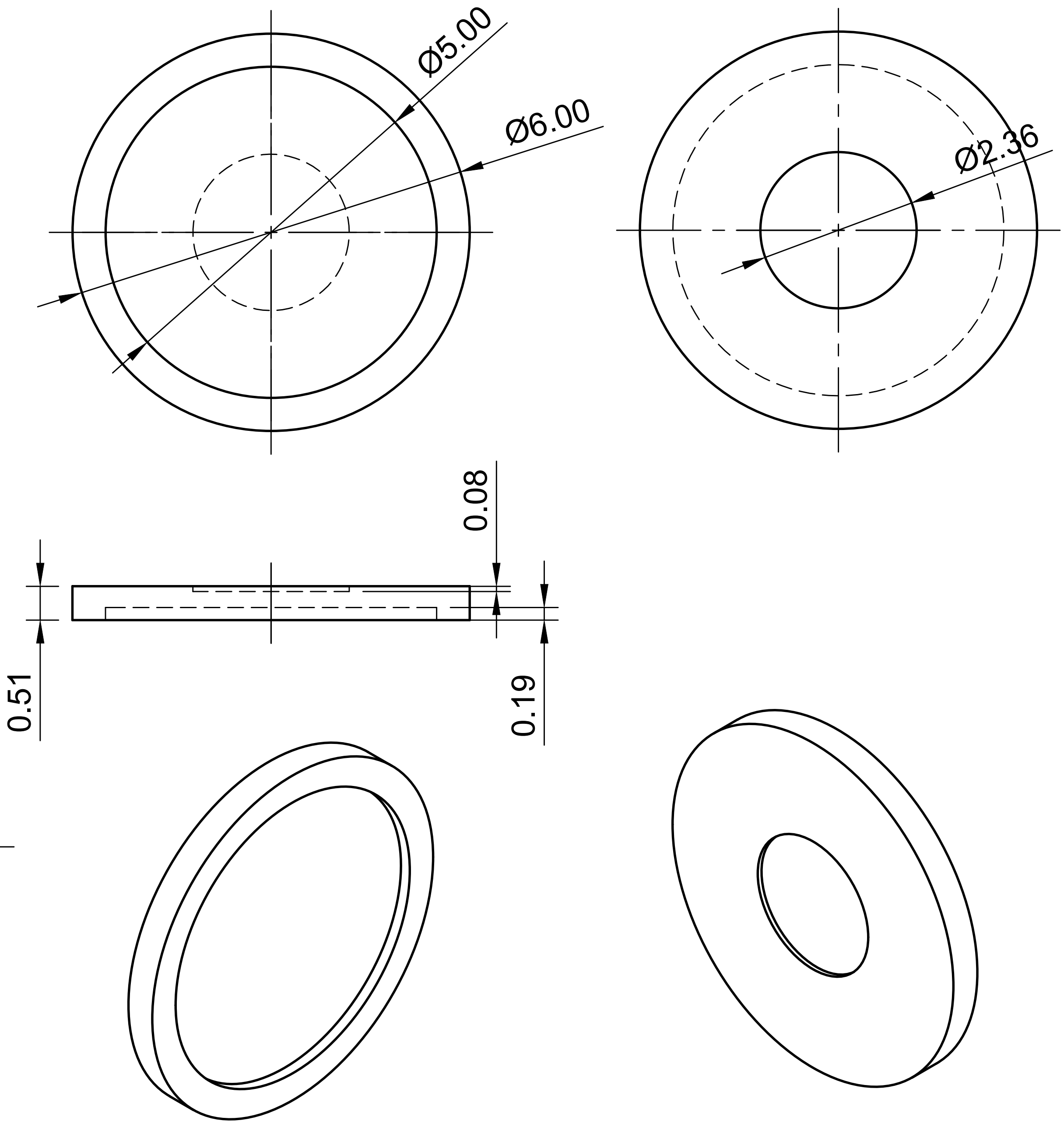

| | | | | |
|---|---|---|---|---|
| Dept. | Technical reference | Created by<br>Miriam Weber 14.04.2026 | Approved by | |
| 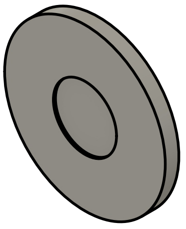 | | Document type | Document status | |
| | | Title<br>Biopsy_Lid_200um_spacer | DWG No. | |
| | | | Rev. / Date of issue | Sheet<br>1/1 |

### Biopsy Lid - 120 µm spacer

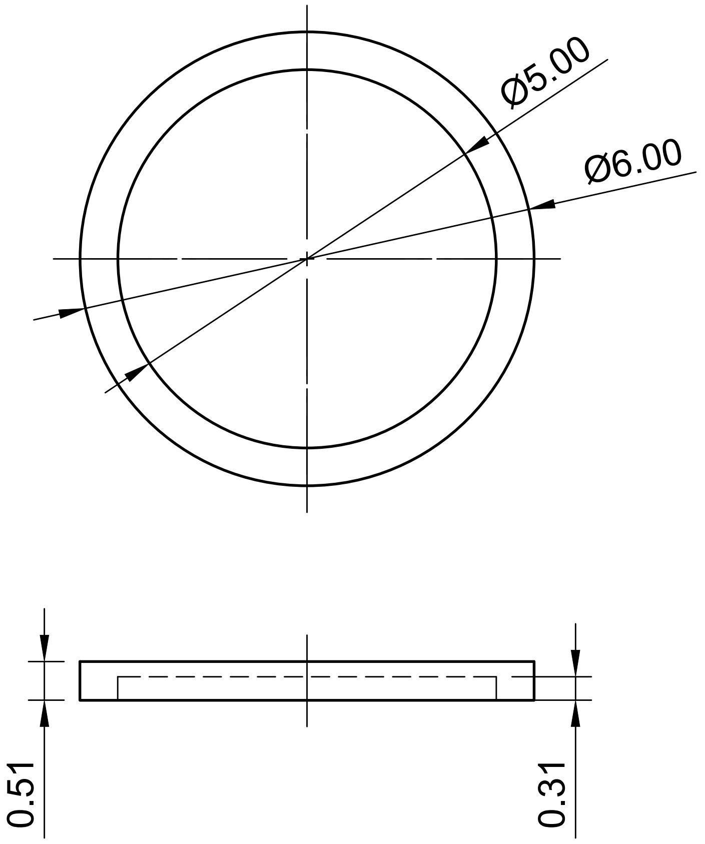

| Dept. | Technical reference | Created by | Approved by |
|---|---|---|---|
| | | Miriam Weber 14.04.2026 | |

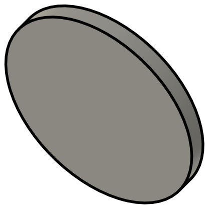

| Document type | Document status |
|---|---|

| Title | DWG No. |
|---|---|
| Biopsy_Flat_Lid_120um_spacer | |

| Rev. | Date of issue | Sheet |
|---|---|---|
| | | 1/1 |

### Biopsy Lid - 60 µm spacer

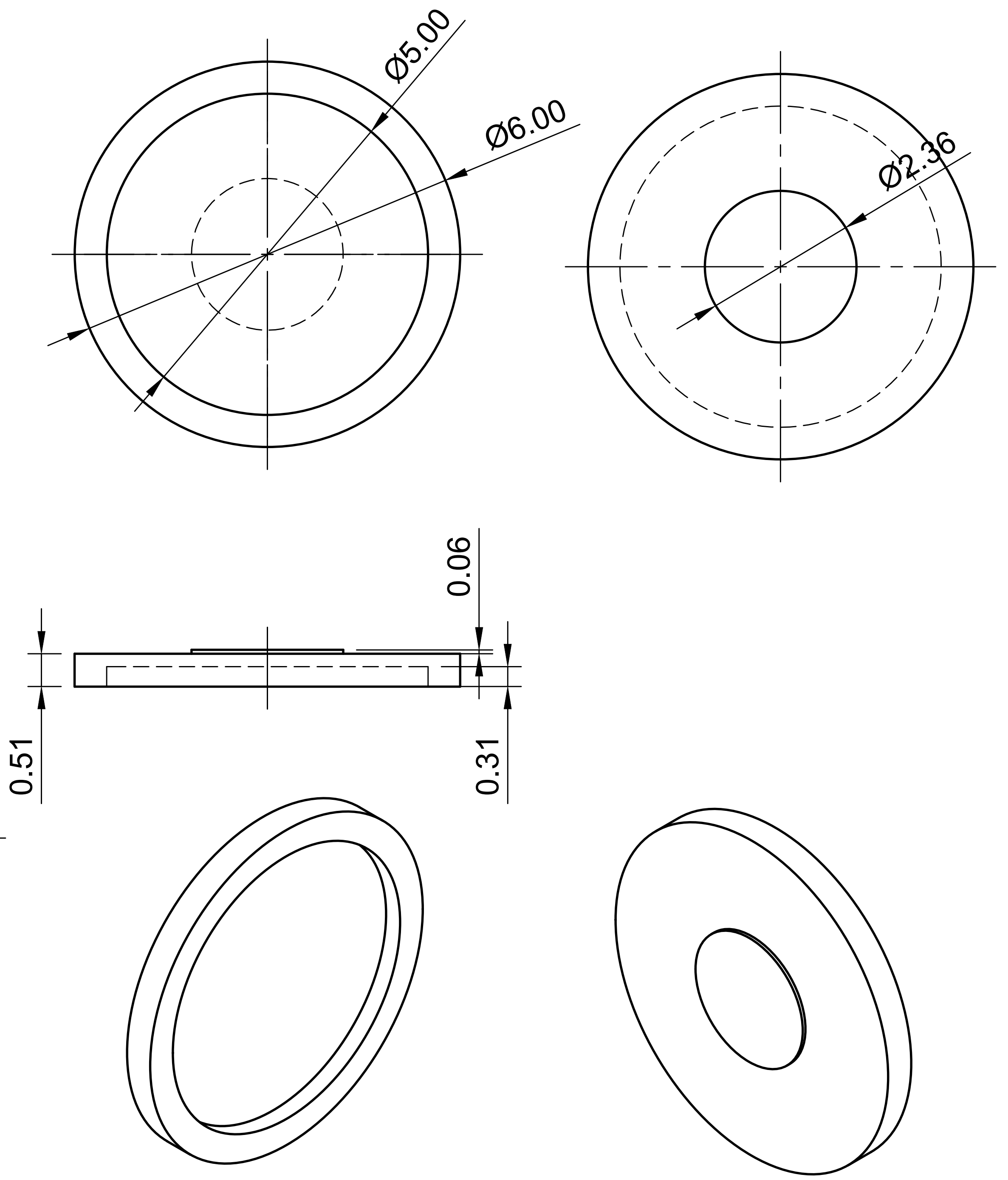

| Dept. | Technical reference | Created by<br>Miriam Weber 14.04.2026 | Approved by | | |
|---|---|---|---|---|---|
| 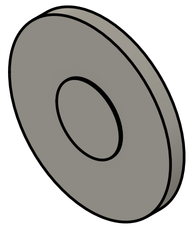 | | Document type | Document status | | |
| | | Title<br>Biopsy_Lid_60um_spacer | DWG No. | | |
| | | | Rev. | Date of issue | Sheet<br>1/1 |
